# A multiscale X-ray phase-contrast tomography dataset of whole human left lung

**DOI:** 10.1101/2021.11.20.469361

**Authors:** R. Patrick Xian, Claire L. Walsh, Stijn E. Verleden, Willi L. Wagner, Alexandre Bellier, Sebastian Marussi, Maximilian Ackermann, Danny D. Jonigk, Joseph Jacob, Peter D. Lee, Paul Tafforeau

**Author notes:** These authors contributed equally to this work (R. Patrick Xian, Paul Tafforeau). corresponding authors: Paul Tafforeau, R. Patrick Xian, Peter D. Lee.

## Abstract

Technological advancements in X-ray imaging using bright and coherent synchrotron sources now allows to decouple sample size and resolution, while maintaining high sensitivity to the microstructure of soft, partially dehydrated tissues. The recently developed imaging technique, hierarchical phase-contrast tomography, is a comprehensive approach to address the challenge of organ-scale (up to tens of centimeters) soft tissue imaging with resolution and sensitivity down to the cellular level. Using this technique, we imaged *ex vivo* an entire human left lung at an isotropic voxel size of 25.08 *μ*m along with local zooms down to 6.05 - 6.5 *μ*m and 2.45 - 2.5 *μ*m in voxel size. The high tissue contrast offered by the fourth-generation synchrotron source at the European Synchrotron Radiation Facility reveals complex multiscale anatomical constitution of the human lung from the macroscopic (centimeter) down to the microscopic (micrometer) scale. The dataset provides complete organ-scale 3D information of the secondary pulmonary lobules and delineates the microstructure of lung nodules with unprecedented detail.

## Background & Summary

The human lung is among the largest solid organs in the human body. Traditionally, studies of lung microanatomy at the organ scale require lengthy operations in targeted sampling, tissue preparation, histological staining and sectioning^1,2^. Nowadays, *ex vivo* clinical evaluations of whole lung microstructures are carried out without sectioning using absorption-contrast micro-CT at around 100 *μ*m voxel size, then a limited area may be selected to image at higher resolution using histology^3–5^. X-ray phase-contrast imaging^6^ provides higher sensitivity and contrast than laboratory micro-CT^7^. Compared with optical virtual histology^8^, X-ray phase contrast from free-space propagation requires no imaging optics and, at the same time, removes the need for laborious tissue clearing and staining that are essential for optical imaging. The compatibility of X-ray phase-contrast imaging with existing X-ray sources facilitates its gradual adoption and transition from preclinical research to clinical diagnostics^9,10^. At synchrotron facilities, systematic upgrades^11,12^ in the X-ray source and imaging techniques over the past decades provide the means to tackle biological questions on meaningful scales and resolution^13–19^. Although synchrotron-based X-ray imaging can access finer anatomical detail than laboratory micro-CT^18,20–22^, many bioimaging scenarios require further upscaling of the imaging throughput and accommodation of large sample size while maintaining microscopic resolution^23,24^. Thanks to the high X-ray photon flux and coherence achieved at modern fourth-generation synchrotron sources and careful design of the imaging protocol, it is now possible to image complete, large, partially dehydrated human organs in their entirety at micrometer resolution using hierarchical phase-contrast tomography (HiP-CT)^25^. It is a single-modality, multiscale imaging technique that employs propagation phase contrast from high-energy, polychromatic X-rays, flat-field correction, attenuation scanning protocol, along with efficient tomographic sampling and stitching pipeline to cover large, soft-tissue organs entirely. The imaging protocol of HiP-CT starts with a two-step tomographic sampling of the whole organ (full-field tomography), followed by progressive zooming in to selective features of the microanatomy through local tomographies at various resolutions compatible with the relevant anatomical context. The imaging technique takes reference from a separate container (reference jar) for flat-field correction to enhance the soft tissue contrast. The organ in the sample jar is embedded in 70% ethanol solution in water and immobilized with agar blocks throughout imaging (see Fig. 1). We provide here the human left lung dataset imaged by HiP-CT at 25.08 *μ*m voxel size (full organ) and at 6.05 - 6.5 *μ*m and 2.45 - 2.5 *μ*m voxel size for various local volumes of interests (VOIs) accomplished by different post-scintillator coupling optics before the detector. The X-ray imaging experiments were carried out at the European Synchrotron Radiation Facility (ESRF) BM05 beamline using the recently upgraded fourth-generation extremely brilliant X-ray source (ESRF-EBS)^26,27^.

**Figure 1.**
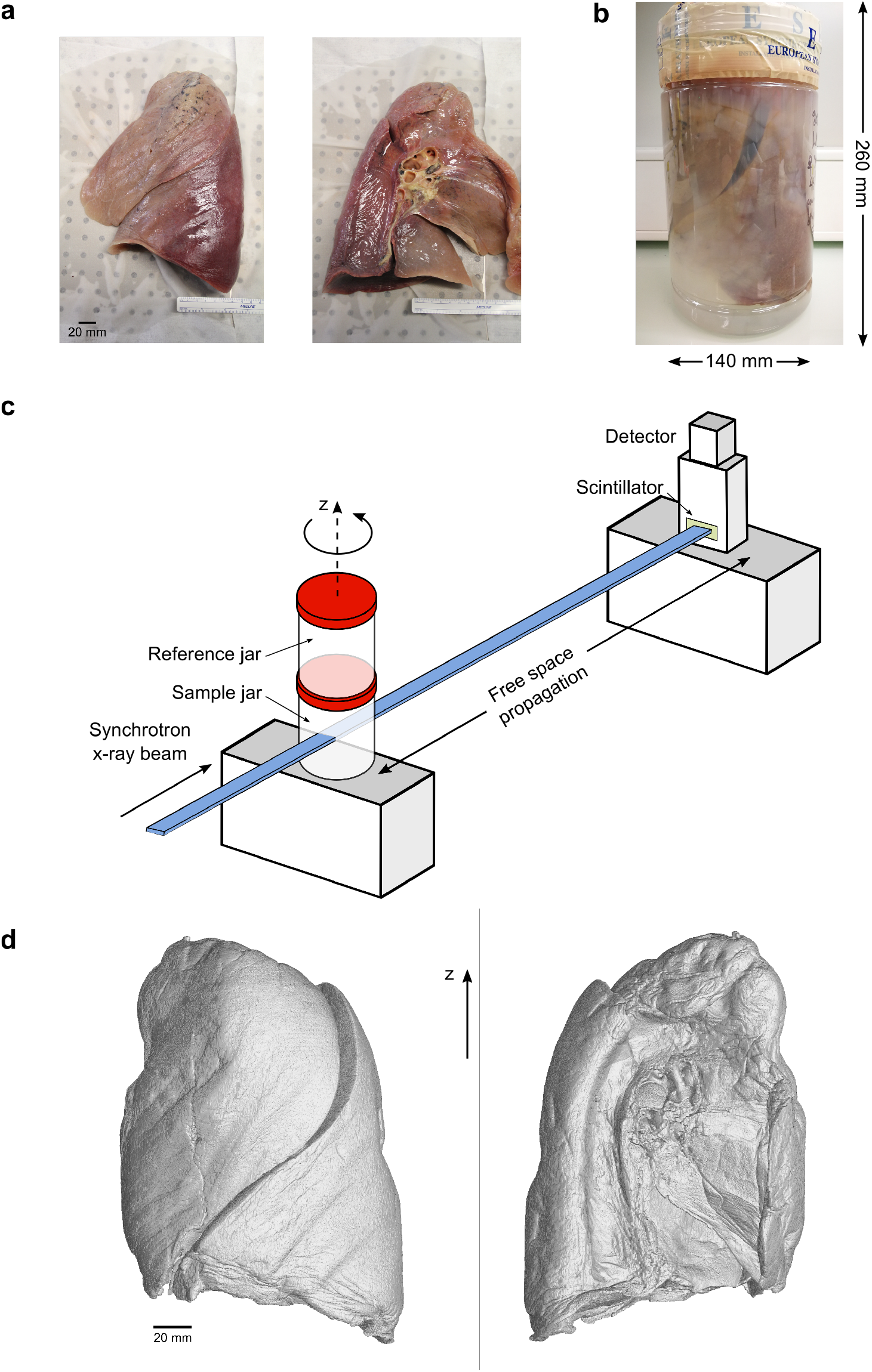
(**a**) A human left lung in lateral (left) and medial (right) view. (**b**) The whole lung is mounted in a sealed, plastic cylindrical jar (140 mm in diameter, 260 mm in height) filled with 70% ethanol solution and agar blocks. (**c**) Sketch of the imaging setup for propagation phase-contrast X-ray tomography at ESRF BM05 beamline. The reference jar contains the same embedding medium as the sample jar. The incident X-ray energy is adjusted to between 70 - 85 keV through filters depending on the resolution requirement. (**d**) Volumetric rendering of the whole left lung imaged at 25.08 *μ*m voxel size using HiP-CT in lateral (left) and medial (right) view.

## Methods

### Lung preparation and mounting

The entire left lung (see Fig. 1a) was harvested from an organ donor, a 94-year-old woman who succumbed to natural causes. Body donation was based on free consent by the donor antemortem. The relevant postmortem medical procedures were carried out at Laboratoire d’Anatomie des Alpes Françaises (LADAF) according to the Quality Appraisal for Cadaveric Studies scale recommendations^28^. All dissections respected the memory of the deceased. The protocols for transport and imaging were approved by the French legislation for body donation. The body of the deceased donor was embalmed and the lung preparations were carried out at ~ 36 hours postmortem. The lung was instilled through the trachea with a 4% formalin solution using 30 cm of water column positive pressure. The trachea was then ligatured to maintain the inflated configuration in order to fix the lungs in a non-collapsed state. The body was then kept at 4 °C for 3 days before the dissection. Once removed, the lungs were immersed in 4% formalin solution for 3 more days. Afterwards, it was successively immersed in ethanol solutions with increasing concentration up to 70% (volume fraction). The lung was kept inflated during ethanol dehydration by repeatedly pushing the solution through its main bronchus with a syringe. The significantly lower density of ethanol (789 kg/m^3^) compared with water (1000 kg/m^3^) provides a high soft tissue base contrast^29,30^.

We used a PET (polyethylene terephthalate) jar of comparable size to the lung for X-ray imaging due to its commercial availability (Medline Scientific, 3600 mL), high radiation tolerance^31^ and optical transparency in assisting sample alignment and assessment of sample condition during imaging. To secure the lung tightly in place and prevent it from touching the container edges on all sides, we prepared agar blocks (~ 1 cm^3^-sized cubes) and stacked them at the bottom of the jar and around the organ to surround and firmly embed the lung. The gaps between the small agar blocks provide the escape routes for residual gas removal. The sample mounting procedure involves alternated filling of the agar-ethanol mixture and gentle vacuum degassing to minimize the existing microbubbles from dissolved air in the solution environment and within the organ, thereby eliminating their interference with imaging. The degassing procedure used a membrane pump to directly pump^32^ above the PET sample jar with the lid open in a sealed vacuum glass dryer. Prior to imaging, the PET jar containing the lung, ethanol solution and agar embedding was placed in a custom-made sample holder to connect to the rotation stage at the synchrotron beamline^25^.

### Synchrotron X-ray imaging and reconstruction

The implementation and capabilities of HiP-CT have been described in detail in a separate publication^25^. Here, we describe the settings used for lung imaging. All X-ray imaging experiments were carried out at the ESRF bending magnet beamline BM05^33^. The polychromatic synchrotron beam produced at the beamline was passed through a set of filters and then directly used for imaging without additional X-ray optics. The voxel size is effectively controlled by the adjustable visible-light imaging optics situated after the LuAG:Ce X-ray scintillator and before the sCMOS light sensor (PCO edge 4.2 CLHS, PCO AG, Germany). Specifically, the imaging optics include the dzoom (“demagnifying zoom”) and zoom lenses, which cover the ranges of 6.5 - 25.5 *μ*m and 1.3 - 6.3 *μ*m, respectively. Because the synchrotron beam size (with usable area 50 mm × 4 mm at BM05) is considerably smaller than the size of the human left lung (container size 260 mm height, up to 140 mm width at the widest), imaging an entire lung at 25.08 *μ*m voxel size requires stitching together multiple subscans. We used the half-acquisition (or half-object acquisition)^34^ method developed at ESRF for imaging the VOIs at 6.5 *μ*m and 2.5 *μ*m in voxel size. For the entire lung, we developed a quarter-acquisition method^25^ that includes the half-acquisition in combination with an annular scan to cover its complete horizontal extent (see Fig. 2).

**Figure 2.**
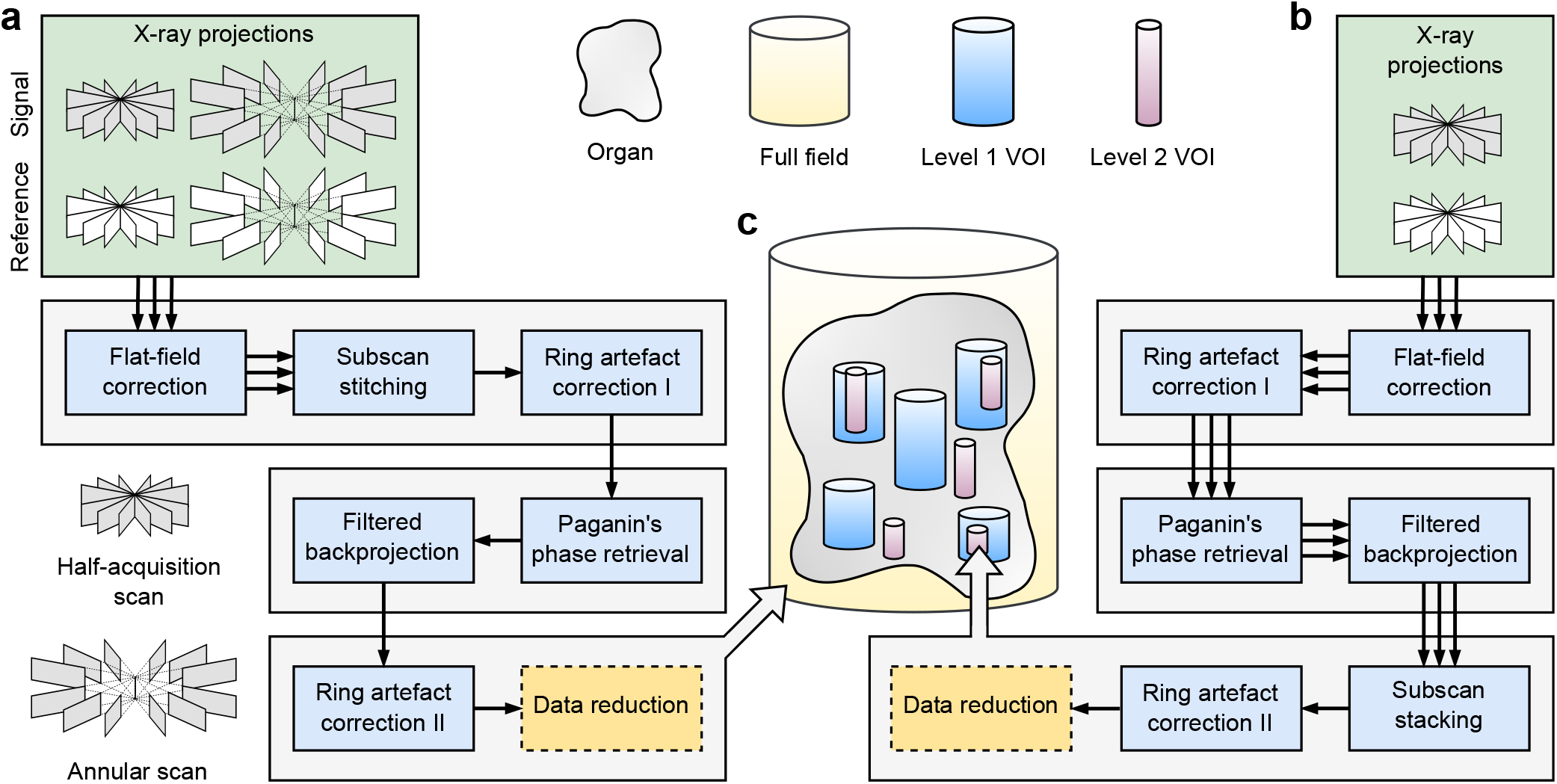
Synchrotron-based hierarchical phase-contrast tomography (HiP-CT) at multiple lengthscales and their associated data acquisition and image reconstruction pipelines. The full-field tomography data at 25.08 *μ*m voxel size are processed with pipeline (**a**). The local tomography data for volumes of interests (VOIs) at 6.05-6.5 *μ*m (level 1) and 2.45 - 2.5 *μ*m (level 2) voxel size are processed with pipeline (**b**). (**c**) A cartoon illustrating the relationships of the various cylindrical volumes imaged with HiP-CT. The triple arrows in the pipeline before merging of the subscans indicate that the same procedure is carried out on each subscan.

Data processing of the measured X-ray projections consists of three stages, pre-reconstruction, reconstruction and post-reconstruction, which are illustrated in separate rows in Fig. 2. Ring artefacts from the detectors are corrected in two steps: (1) Before reconstruction, the mean of the projections is subtracted from the projections to remove the rings with constant intensity rings; (2) After reconstruction, the residual inhomogeneous intensity rings were removed using the polar transform combined with linear motion blurring filter^35^. Tomographic reconstruction employs the phase and amplitude estimates obtained from Paganin’s method^36^, followed by a 2D unsharp mask of the retrieved phase maps as input for the filtered backprojection algorithm. These reconstruction steps are implemented in PyHST2^37^. Eventually, the processed volumes are converted to 16 bit and binned further to produce the datasets described in Tables 1–2. The reconstruction and postprocessing steps are illustrated for the three types of imaged volumes, respectively, in Fig. 2. We summarize below the imaging and reconstruction protocols for the human lung at each imaged resolution including the key parameters.

- Full-field tomography (whole organ at 25.08 *μ*m voxel size, see Fig. 2a,c): The incident X-ray energy averaged at ~ 85 keV after filters, the propagation distance is 3475 mm. In total, two sets of 9990 projections were measured by the quarter-acquisition method^25^ with an offset of 800 pixels for the half-acquisition. A step size of 2.2 mm in the vertical (z) direction is used to cover the height of the sample jar with a total of 98 quarter-acquisition subscans. Radiograph stitching is carried out to recover a half-acquisition scan^34^ before the reconstruction.
- Local tomography of level 1 VOI (6.5 *μ*m and 6.05 *μ*m voxel size, see Fig. 2b,c): The incident X-ray energy averaged at ~ 88 keV (~ 89 keV) after filters, the propagation distance is 3500 mm (3475 mm) for the VOIs with 6.5 *μ*m (6.05 *μ*m) voxel size. In total, 6000 projections were measured by the half-acquisition method with an offset of 900 pixels. A step size of 2.2 mm in the vertical direction is used to cover the height of the VOIs.
- Local tomography of level 2 VOI (2.5 *μ*m and 2.45 *μ*m voxel size, see Fig. 2b,c): The incident X-ray energy averaged at ~ 77 keV (~ 79 keV), the propagation distance is 1440 mm (1500 mm) for the VOIs with 2.5 *μ*m (2.45 *μ*m) voxel size. In total, 6000 projections were measured by the half-acquisition method with an offset of 900 pixels. A step size of 1.5 mm in the vertical direction is used to cover the height of the VOIs.

**Table 1.** Volumes of interests and their anatomical references to the human left lung sample.

| VOI reference | Height (mm) | Diameter (mm) | Displacement (mm) | Z rotation | Anatomical reference |
| --- | --- | --- | --- | --- | --- |
| $6.5 \mu\text{m}$, VOI1 | 48.79 | 25.06 | (23.3, -9.4, -81.2) | $24.6^\circ$ | upper lobe, apical region |
| $6.5 \mu\text{m}$, VOI2b | 26.74 | 25.05 | (5.8, -15.0, -26.5) | $24.6^\circ$ | upper lobe, medial region |
| $6.5 \mu\text{m}$, VOI3b | 9.11 | 25.04 | (5.5, 0.8, -15.2) | $24.6^\circ$ | interlobar fissure |
| $6.5 \mu\text{m}$, VOI4b | 106.07 | 25.06 | (-20.6, 8.1, 37.1) | $24.6^\circ$ | lower lobe, basal to medial region |
| $6.5 \mu\text{m}$, VOI5 | 59.81 | 25.04 | (-3.3, -15.1, -71.0) | $24.6^\circ$ | upper lobe, apical region |
| $6.05 \mu\text{m}$, VOI1 | 7.61 | 23.12 | (-12.5, -23.5, -67.9) | $2^\circ$ | upper lobe, apical region |
| $6.05 \mu\text{m}$, VOI2 | 42.17 | 23.10 | (-7.0, 11.4, 65.5) | $0^\circ$ | lower lobe, basal region |
| $6.05 \mu\text{m}$, VOI6 | 48.85 | 23.09 | (-19.3, -4.9, 59.7) | $0^\circ$ | lower lobe, basal region |
| $2.5 \mu\text{m}$, VOI1 | 32.00 | 9.69 | (18.4, -19.5, -84.5) | $24.6^\circ$ | upper lobe, apical region |
| $2.5 \mu\text{m}$, VOI2b | 14.00 | 9.70 | (0.3, -20.7, -83.6) | $24.6^\circ$ | upper lobe, apical region |
| $2.5 \mu\text{m}$, VOI3 | 14.00 | 9.69 | (-12.6, -24.6, -65.7) | $24.6^\circ$ | upper lobe, apical region |
| $2.5 \mu\text{m}$, VOI4 | 14.00 | 9.69 | (20.5, -19.0, -43.0) | $24.6^\circ$ | upper lobe, medial region |
| $2.5 \mu\text{m}$, VOI5 | 38.25 | 9.67 | (-22.1, 6.4, 63.5) | $24.6^\circ$ | lower lobe, basal region |
| $2.45 \mu\text{m}$, VOI1 | 8.52 | 9.33 | (-12.6, -23.7, -67.5) | $4.8^\circ$ | upper lobe, apical region |
| $2.45 \mu\text{m}$, VOI2 | 11.93 | 9.33 | (-7.7, 12.0, 63.0) | $2^\circ$ | lower lobe, basal region |
| $2.45 \mu\text{m}$, VOI6 | 4.90 | 9.33 | (-19.3, -4.9, 65.1) | $0^\circ$ | lower lobe, basal region |

**Table 2.**
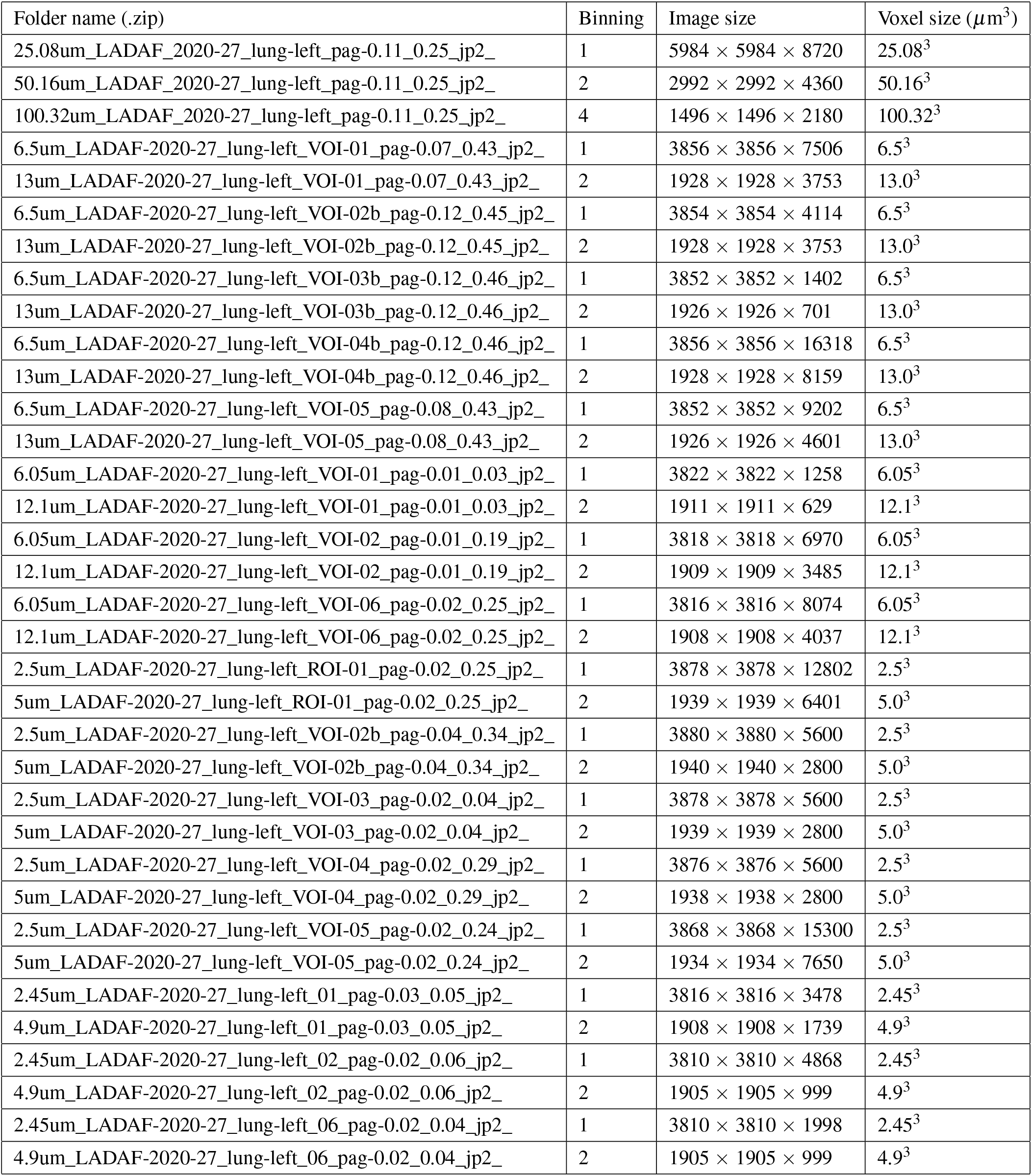
Details of the hierarchical X-ray phase-contrast tomography data for human left lung.

### Volume selection and anatomical reference

Besides the full-field tomography of the entire lung, subsequent smaller VOIs were selected with representative features and imaged with local tomography at higher resolution, including 6.5 *μ*m (5 locations) and 6.05 *μ*m (3 locations) for level 1 and 2.5 *μ*m (5 locations) and 2.45 *μ*m (3 locations) for level 2 VOIs, respectively. All VOIs have a cylindrical field of view about the rotation axis after removing the boundary artefacts from local tomographic reconstruction. To obtain the displacements and rotations, the VOIs are spatially registered to the whole lung data by hand in VGStudio Max (version 3.4) and the procedure to apply them is described in Usage Notes. The sizes of the VOIs, their displacements and rotations with respect to the center of the whole lung data are listed in Table 1 and illustrated in Fig. 3a-d. In addition, we provide brief anatomical references to the VOI spatial locations in Table 1 with respect to the whole lung data at 25.08 *μ*m. To retain traceable data provenance, we keep the same alphanumeric label of the VOIs as used in the original experiments. Fig. 3e visualizes two selected VOIs in the lower lobe of the lung.

**Figure 3.**
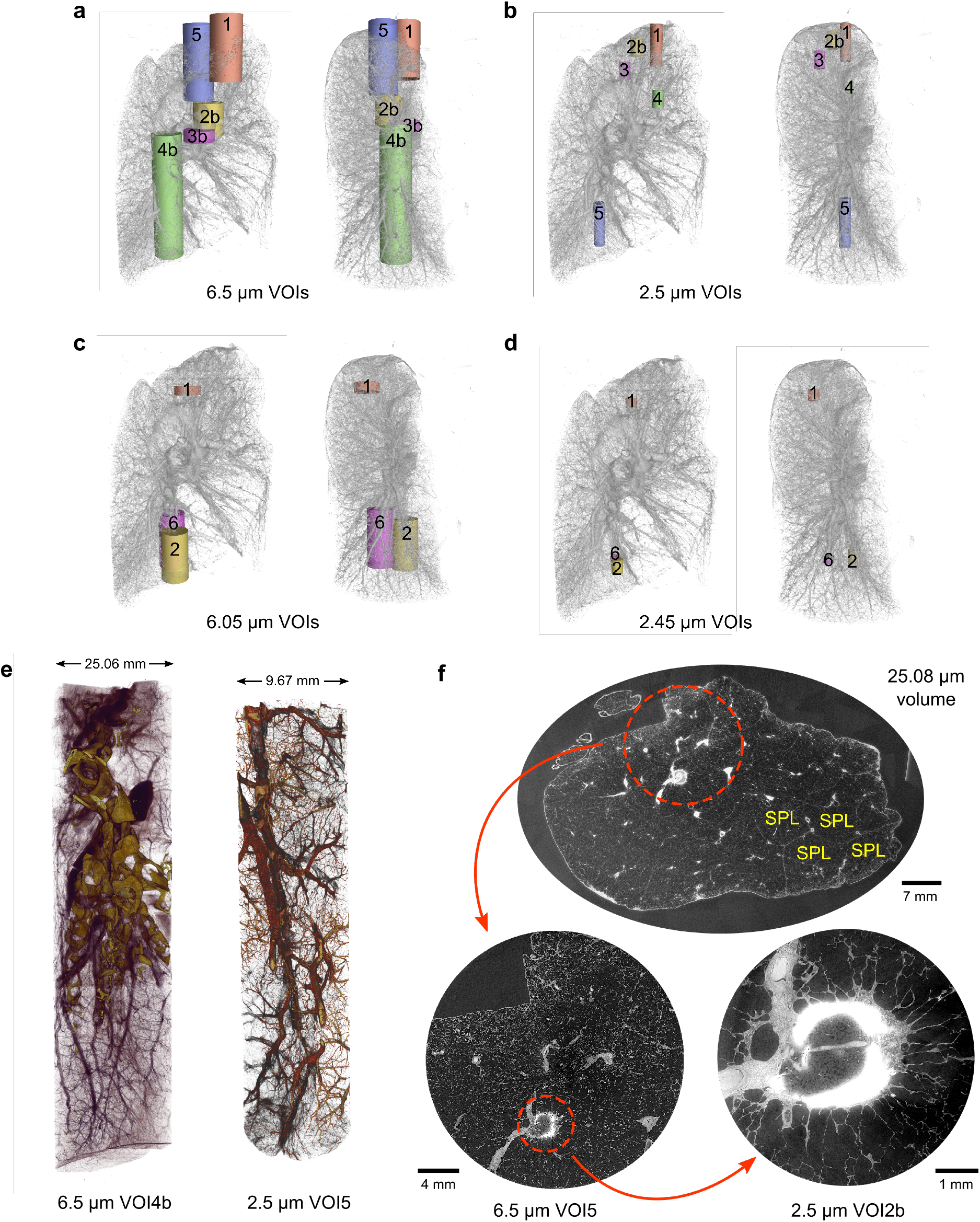
Exploration of the HiP-CT dataset of the human left lung. (**a**-**d**) Spatial correspondences of the measured cylindrical VOIs at different resolutions within the entire left lung. For each set of VOIs, both the medial (left) and sagittal (right) views are shown. The VOI label corresponds to the assignment in Table 1. (**e**) Renderings of two imaged VOIs with 6.5 *μ*m and 2.5 *μ*m voxel sizes, respectively. (**f**) From the whole lung and local zoom data, we visualize the anatomical detail of a spiculated lung nodule in the apical region of the lung on multiple lengthscales. The interlobular septa and perilobular vasculature of the secondary pulmonary lobules (SPLs) are clearly visible.

## Data Records

We provide the volumetric data after reconstruction and post-processing as greyscale (16 bit) 2D image slices in JPEG2000 format stored in zipped folders. The compression level of JPEG2000 is carefully chosen to ensure minimal difference from the original TIFF-formatted data when they are used for feature quantification or image segmentation. We list the details of the deposited data is in Table 2. All data have been deposited at an ESRF data repository (https://human-organ-atlas.esrf.eu/) with digital object identifiers (DOIs) assigned to each scanned volume as listed in Table 3. Each DOI refers to a volume at full resolution and its binned versions. For all volumes of interests measured by local tomography, both the full resolution data (Binning = 1) and the 2× binned version (Binning = 2) are provided, while for the whole lung data, the 4× binned version (Binning = 4) is also provided. The metadata information in Table 1 is also provided in the corresponding text file contained in each data deposit.

**Table 3.** Information about the data records.

| Data description | Index | DOI |
| --- | --- | --- |
| Full-field tomography data at 25.08 $\mu\text{m}$ voxel size and binned versions (50.16 $\mu\text{m}$, 100.32 $\mu\text{m}$ voxel sizes) | | [10.15151/ESRF-DC-572196058](https://doi.org/10.15151/ESRF-DC-572196058) |
| Local tomography data at 6.5 $\mu\text{m}$ voxel size and binned version (13.0 $\mu\text{m}$ voxel size) | VOI1 | [10.15151/ESRF-DC-572235698](https://doi.org/10.15151/ESRF-DC-572235698) |
|  | VOI2b | [10.15151/ESRF-DC-572236926](https://doi.org/10.15151/ESRF-DC-572236926) |
|  | VOI3b | [10.15151/ESRF-DC-572237999](https://doi.org/10.15151/ESRF-DC-572237999) |
|  | VOI4b | [10.15151/ESRF-DC-572240585](https://doi.org/10.15151/ESRF-DC-572240585) |
|  | VOI5 | [10.15151/ESRF-DC-572242236](https://doi.org/10.15151/ESRF-DC-572242236) |
| Local tomography data at 6.05 $\mu\text{m}$ voxel size and binned version (12.1 $\mu\text{m}$ voxel size) | VOI1 | [10.15151/ESRF-DC-572230985](https://doi.org/10.15151/ESRF-DC-572230985) |
|  | VOI2 | [10.15151/ESRF-DC-572231249](https://doi.org/10.15151/ESRF-DC-572231249) |
|  | VOI6 | [10.15151/ESRF-DC-572232527](https://doi.org/10.15151/ESRF-DC-572232527) |
| Local tomography data at 2.5 $\mu\text{m}$ voxel size and binned version (5.0 $\mu\text{m}$ voxel size) | VOI1 | [10.15151/ESRF-DC-572221364](https://doi.org/10.15151/ESRF-DC-572221364) |
|  | VOI2b | [10.15151/ESRF-DC-572222783](https://doi.org/10.15151/ESRF-DC-572222783) |
|  | VOI3 | [10.15151/ESRF-DC-572222987](https://doi.org/10.15151/ESRF-DC-572222987) |
|  | VOI4 | [10.15151/ESRF-DC-572229061](https://doi.org/10.15151/ESRF-DC-572229061) |
|  | VOI5 | [10.15151/ESRF-DC-572229315](https://doi.org/10.15151/ESRF-DC-572229315) |
| Local tomography data at 2.45 $\mu\text{m}$ voxel size and binned version (4.9 $\mu\text{m}$ voxel size) | VOI1 | [10.15151/ESRF-DC-572191396](https://doi.org/10.15151/ESRF-DC-572191396) |
|  | VOI2 | [10.15151/ESRF-DC-572191782](https://doi.org/10.15151/ESRF-DC-572191782) |
|  | VOI6 | [10.15151/ESRF-DC-572194514](https://doi.org/10.15151/ESRF-DC-572194514) |

## Technical Validation

Although the radiation dose during tomographic scans is well below the tissue damage threshold^25^, due to radiation-induced bubble formation, the sample went through re-degassing before further measurements are made. A consequence is that not all of the VOIs have been imaged consecutively during the same beamtime. In the course of re-degassing, the sample was kept in the container to maintain its position. The jar is then placed into the synchrotron X-ray beamline for further imaging. Care is exercised in the process such that the VOIs scanned before and after re-degassing can be registered to the whole volume without large deformation.

In the imaged volumes, contrast is produced by the local density differences between the lung tissue constituents and the hollow structures (e.g. airway, alveoli, blood vessels) filled with ethanol solution (see Fig. 3e-f). Within the whole lung data at a voxel size of 25.08 *μ*m, the interlobular septa, the boundaries of the secondary pulmonary lobules^38,39^ and the perilobular vasculature, are clearly visible (see Fig. 3f). At high spatial resolution, the local density difference increasingly becomes the dominant contributor to image contrast for VOIs^25^. The consistent contrast across lengthscales provides unprecedentedly detailed information for the study of lung morphology in health and disease.

## Usage Notes

The multiscale healthy human lung data presented here have been used as clinical control data in studies comparing damage within the lung microstructure due to Covid-19 infection^25^. The datasets are deposited as 2D image slices perpendicular to the rotation axis (z in Fig. 1) in tomography geometry. These images may be directly loaded into any typical image processing software for viewing or further quantification. To align the VOIs to the whole lung data, the following transform should be applied to the VOI,

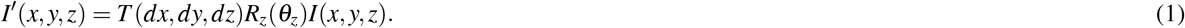

Here *I* and *I*′ are intensity-valued volumetric data, *T* is the 3D translation operator, and *R_z_* the 3D rotation operator around *z* axis (see Fig. 1b-c). The displacement vector (*dx, dy, dz*) and *z* rotation angle *θ_z_* for each VOI is listed in Table 1. The greyscale ranges of the images are selected with an intensity margin to avoid saturation. Viewing directly by eye may require threshold adjustment.

## Code availability

The code used for the preprocessing, tomographic reconstruction and postprocessing is available on GitHub (https://github.com/HiPCTProject/Tomo_Recon).

## Acknowledgements

We thank S. Bayat (ISERM), P. Masson (LADAF) for extracting the lung from the body donor, H. Reichert (ESRF) and R. Torii (UCL) for general support of the project, and C. Muzelle, R. Homs, C. Jarnias, F. Cianciosi, P. Vieux, P. Cook, L. Capasso and A. Mirone for their help in the X-ray imaging setup developments and improvements. We thank A. de Maria, A. Bocciarelli, M. Bodin, J.-F. Perrin, and A. Goetz at ESRF for support on the Human Organ Atlas database. The authors also thank C. Werlein, R. Engelhardt, A. M. Brechlin, C. Petzold and N. Kroenke. This project has been made possible in part by grants number 2020-225394 from the Chan Zuckerberg Initiative DAF, an advised fund of Silicon Valley Community Foundation, The ESRF - funding proposal md1252, the Royal Academy of Engineering (PDL - CiET1819/10) and the MRC (MR/R025673/1). M.A. acknowledges grants from the National Institutes of Health (HL94567 and HL134229). D.D.J. acknowledges the European Consolidator Grant, XHale (ref. no. 771883). J.J. acknowledges Wellcome Trust Clinical Research Career Development Fellowship 209553/Z/17/Z and the National Institute for Health Research University College London Hospital Biomedical Research Centre.

## Author contributions statement

P.D.L. and P.T. conceived the experiment. P.D.L., C.L.W. and P.T. coordinated the collaboration. A.B. harvested the lung from the organ donor and prepared the lung for imaging along with P.T.. S.M. designed the sample holder for organ imaging. P.T. conducted the imaging experiment at ESRF BM05 beamline and reconstructed the volumetric data. R.P.X. analyzed the data with help and instructions from P.T., S.V., W.L.W., J.J., M.A. and D.D.J.. R.P.X. wrote the first version of the manuscript. All authors reviewed and discussed the manuscript to bring it to the final form.

## References

1. Weibel, E. R. Morphometry of the Human Lung (Springer Berlin Heidelberg, Berlin, Heidelberg, 1963).

2. Ochs, M. et al. The Number of Alveoli in the Human Lung. Am. J. Respir. Critical Care Medicine 169, 120–124, 10.1164/rccm.200308-1107OC (2004).

3. Katsamenis, O. L. et al. X-ray Micro-Computed Tomography for Nondestructive Three-Dimensional (3D) X-ray Histology. The Am. J. Pathol. 189, 1608–1620, 10.1016/j.ajpath.2019.05.004 (2019).

4. Vasilescu, D. M. et al. Comprehensive stereological assessment of the human lung using multiresolution computed tomography. J. Appl. Physiol. 128, 1604–1616, 10.1152/japplphysiol.00803.2019 (2020).

5. Verleden, S. E. et al. Small airway loss in the physiologically ageing lung: a cross-sectional study in unused donor lungs. The Lancet Respir. Medicine 9, 167–174, 10.1016/S2213-2600(20)30324-6 (2021).

6. Endrizzi, M. X-ray phase-contrast imaging. Nucl. Instruments Methods Phys. Res. Sect. A: Accel. Spectrometers, Detect. Assoc. Equip. 878, 88–98, 10.1016/j.nima.2017.07.036 (2018).

7. Ritman, E. L. Current Status of Developments and Applications of Micro-CT. Annu. Rev. Biomed. Eng. 13, 531–552, 10.1146/annurev-bioeng-071910-124717 (2011).

8. Liu, J. T. C. et al. Harnessing non-destructive 3D pathology. Nat. Biomed. Eng. 5, 203–218, 10.1038/s41551-020-00681-x (2021).

9. Bravin, A., Coan, P. & Suortti, P. X-ray phase-contrast imaging: from pre-clinical applications towards clinics. Phys. Medicine Biol. 58, R1–R35, 10.1088/0031-9155/58/1/R1 (2013).

10. Momose, A. X-ray phase imaging reaching clinical uses. Phys. Medica 79, 93–102, 10.1016/j.ejmp.2020.11.003 (2020).

11. Bilderback, D. H., Elleaume, P. & Weckert, E. Review of third and next generation synchrotron light sources. J. Phys. B: At. Mol. Opt. Phys. 38, S773–S797, 10.1088/0953-4075/38/9/022 (2005).

12. Weckert, E. The potential of future light sources to explore the structure and function of matter. IUCrJ 2, 230–245, 10.1107/S2052252514024269 (2015).

13. Wagner, W. L. et al. Towards synchrotron phase-contrast lung imaging in patients – a proof-of-concept study on porcine lungs in a human-scale chest phantom. J. Synchrotron Radiat. 25, 1827–1832, 10.1107/S1600577518013401 (2018).

14. Ding, Y. et al. Computational 3D histological phenotyping of whole zebrafish by X-ray histotomography. eLife 8, 10.7554/eLife.44898 (2019).

15. Zdora, M.-C. et al. X-ray phase tomography with near-field speckles for three-dimensional virtual histology. Optica 7, 1221, 10.1364/OPTICA.399421 (2020).

16. Kuan, A. T. et al. Dense neuronal reconstruction through X-ray holographic nano-tomography. Nat. Neurosci. 23, 1637–1643, 10.1038/s41593-020-0704-9 (2020).

17. Longo, E. et al. X-ray Zernike phase contrast tomography: 3D ROI visualization of mm-sized mice organ tissues down to sub-cellular components. Biomed. Opt. Express 11, 5506, 10.1364/BOE.396695 (2020).

18. Umetani, K., Okamoto, T., Saito, K., Kawata, Y. & Niki, N. 36M-pixel synchrotron radiation micro-CT for whole secondary pulmonary lobule visualization from a large human lung specimen. Eur. J. Radiol. Open 7, 100262, 10.1016/j.ejro.2020.100262 (2020).

19. Westöö, C. et al. Distinct types of plexiform lesions identified by synchrotron-based phase-contrast micro-CT. Am. J. Physiol. Cell. Mol. Physiol. 321, L17–L28, 10.1152/ajplung.00432.2020 (2021).

20. Litzlbauer, H. D. et al. Synchrotron-Based Micro-CT Imaging of the Human Lung Acinus. The Anat. Rec. Adv. Integr. Anat. Evol. Biol. 293, 1607–1614, 10.1002/ar.21161 (2010).

21. Elfarnawany, M. et al. Micro-CT versus synchrotron radiation phase contrast imaging of human cochlea. J. Microsc. 265, 349–357, 10.1111/jmi.12507 (2017).

22. Norvik, C. et al. Synchrotron-based phase-contrast micro-CT as a tool for understanding pulmonary vascular pathobiology and the 3-D microanatomy of alveolar capillary dysplasia. Am. J. Physiol. Cell. Mol. Physiol. 318, L65–L75, 10.1152/ajplung.00103.2019 (2020).

23. Cunningham, J. A., Rahman, I. A., Lautenschlager, S., Rayfield, E. J. & Donoghue, P. C. A virtual world of paleontology. Trends Ecol. & Evol. 29, 347–357, 10.1016/j.tree.2014.04.004 (2014).

24. Du, M. et al. Upscaling X-ray nanoimaging to macroscopic specimens. J. Appl. Crystallogr. 54, 386–401, 10.1107/S1600576721000194 (2021).

25. Walsh, C. L. et al. Imaging intact human organs with local resolution of cellular structures using hierarchical phase-contrast tomography. Nat. Methods 1–10, 10.1038/s41592-021-01317-x (2021).

26. Raimondi, P. ESRF-EBS: The Extremely Brilliant Source Project. Synchrotron Radiat. News 29, 8–15, 10.1080/08940886.2016.1244462 (2016).

27. Rack, A. Hard X-ray Imaging at ESRF: Exploiting Contrast and Coherence with the New EBS Storage Ring. Synchrotron Radiat. News 33, 20–28, 10.1080/08940886.2020.1751519 (2020).

28. Wilke, J. et al. Appraising the methodological quality of cadaveric studies: validation of the QUACS scale. J. Anat. 226, 440–446, 10.1111/joa.12292 (2015).

29. Shirai, R. et al. Enhanced renal image contrast by ethanol fixation in phase-contrast X-ray computed tomography. J. Synchrotron Radiat. 21, 795–800, 10.1107/S1600577514010558 (2014).

30. Patzelt, M. et al. Ethanol fixation method for heart and lung imaging in micro-CT. Jpn. J. Radiol. 37, 500–510, 10.1007/s11604-019-00830-6 (2019).

31. Gamma Radiation Resistance: Definition & Values For Plastics. https://omnexus.specialchem.com/polymer-properties/properties/gamma-radiation-resistance.

32. Luyckx, G. & Ceulemans, J. Deoxygenation, Deaeration and Degassing: A Survey and Evaluation of Methods. Bull. des Sociétés Chimiques Belges 96, 151–163, 10.1002/bscb.19870960214 (1987).

33. Ziegler, E. et al. The ESRF BM05 Metrology Beamline: Instrumentation And Performance Upgrade. In AIP Conference Proceedings, vol. 705, 436–439, 10.1063/1.1757827 (AIP, 2004).

34. Kyrieleis, A., Ibison, M., Titarenko, V. & Withers, P. Image stitching strategies for tomographic imaging of large objects at high resolution at synchrotron sources. Nucl. Instruments Methods Phys. Res. Sect. A: Accel. Spectrometers, Detect. Assoc. Equip. 607, 677–684, 10.1016/j.nima.2009.06.030 (2009).

35. Lyckegaard, A., Johnson, G. & Tafforeau, P. Correction of ring artifacts in X-ray tomographic images. Int. J. Tomogr. Stat. 18, 1–9 (2011).

36. Paganin, D., Mayo, S. C., Gureyev, T. E., Miller, P. R. & Wilkins, S. W. Simultaneous phase and amplitude extraction from a single defocused image of a homogeneous object. J. Microsc. 206, 33–40, 10.1046/j.1365-2818.2002.01010.x (2002).

37. Mirone, A., Brun, E., Gouillart, E., Tafforeau, P. & Kieffer, J. The PyHST2 hybrid distributed code for high speed tomographic reconstruction with iterative reconstruction and a priori knowledge capabilities. Nucl. Instruments Methods Phys. Res. Sect. B: Beam Interactions with Mater. Atoms 324, 41–48, 10.1016/j.nimb.2013.09.030 (2014).

38. Bergin, C., Roggli, V., Coblentz, C. & Chiles, C. The secondary pulmonary lobule: normal and abnormal CT appearances. Am. J. Roentgenol. 151, 21–25, 10.2214/ajr.151.1.21 (1988).

39. Webb, W. R. Thin-Section CT of the Secondary Pulmonary Lobule: Anatomy and the Image—The 2004 Fleischner Lecture. Radiology 239, 322–338, 10.1148/radiol.2392041968 (2006).

